# My voice therefore I spoke: sense of agency over speech enhanced in hearing self-voice

**DOI:** 10.1101/2020.11.20.392308

**Authors:** Ryu Ohata, Tomohisa Asai, Shu Imaizumi, Hiroshi Imamizu

**Affiliations:** Department of Psychology, Graduate School of Humanities and Sociology, The University of Tokyo, Hongo 7-3-1, Bunkyo-ku, Tokyo 113-0033, Japan; Cognitive Mechanisms Laboratories, Advanced Telecommunications Research Institute International (ATR), Keihanna Science City, Kyoto 619-0288, Japan; Institute for Education and Human Development, Ochanomizu University, Otsuka 2-1-1, Bunkyo-ku, Tokyo 112-8610, Japan; Research into Artifacts, Center for Engineering, The University of Tokyo, Hongo 7-3-1, Bunkyo-ku, Tokyo 113-0033, Japan

## Abstract

The subjective experience of causing an action is known as the sense of agency. Dysfunction in the sense of agency has been suggested as a cause of auditory hallucinations (AHs), an important diagnostic criterion for schizophrenia. However, agency over speech has not been extensively characterized in previous empirical studies. Here, we examine both implicit and explicit measures of the sense of agency and reveal bottom-up and top-down components that constitute self-agency during speech. The first is action-outcome causality, which is perceived based on a low-level sensorimotor process when hearing their own voice following their speech. The second component is self-voice identity, which is embedded in the acoustic quality of voice and dominantly influences agency over speech at the cognitive judgment level. Our findings provide profound insight into the sense of agency over speech and present an informative perspective for understanding aberrant experience in AHs.

## Introduction

The subjective experience that “I” am the one who is causing an action, referred to as the sense of agency, is the fundamental aspect of the sense of self (Gallagher, 2000; Haggard, 2017). Although people are not usually aware of its existence, pathological conditions make it evident that this sense is essential in our daily activities. For example, the delusion of control, which is one of the important diagnostic criteria for schizophrenia, denotes an abnormal experience in which an external force controls one’s actions, thoughts, or feelings. Theoretical studies hypothesized that these symptoms arise from dysfunction in the sense of agency (Blakemore et al., 2003; Frith et al., 2000a; Frith et al., 2000b; Gallagher & Trigg, 2016). Psychological experiments have provided supportive evidence of this hypothesis by examining awareness in patients with schizophrenia during hand/limb movements (Haggard et al., 2003; Synofzik et al., 2009; Voss et al., 2017; Voss et al., 2010; Werner et al., 2014). Furthermore, it has been suggested that an auditory hallucination (AH), which is also a representative symptom of schizophrenia, also results from dysfunction in the sense of agency over speech (Blakemore, et al., 2000a; Ford, 2016; Frith, 1992). However, compared with the sense of agency over hand/limb movement, the mechanism underlying the agency over speech has not been extensively characterized.

The theoretical model for the sense of agency, known as the comparator model (Blakemore, et al., 2000b; Frith et al., 2000a), emphasizes the role of the sensorimotor system based on a computational model of motor control. Here, an internal forward model (Miall & Wolpert, 1996) predicts sensory outcomes of action from an efference copy of a motor command and then compares the predicted and actual outcomes in the brain. If the two outcomes are congruent, people perceive causality between their action and consecutive sensory outcomes and, as a result, feel their agency over the action (blue frame in Figure 1). Neurophysiological studies in humans have captured this comparison process in the sensorimotor system for speech. Auditory event-related potential (N1-component) was more greatly suppressed when an undistorted self-voice was fed back to healthy participants compared to a pitch-distorted voice (Heinks-Maldonado et al., 2005). This suppression was suggested as the instantiation of the result of the comparison. Importantly, the suppression did not appear in patients with AHs, which implies that dysfunction of the comparator system in speech is associated with AHs (Ford et al., 2007; Ford & Mathalon, 2005; Heinks-Maldonado et al., 2007; Whitford et al., 2011). Previous studies have, taken together, demonstrated the importance of the predictive sensorimotor system in understanding the mechanism of AHs (Jones & Fernyhough, 2007; Seal et al., 2004). However, to our knowledge, no study has directly investigated whether such a sensorimotor system for speech enables people to perceive causality between their speech and their own voice, which would be necessary for the sense of agency during speech.

**Figure 1.**
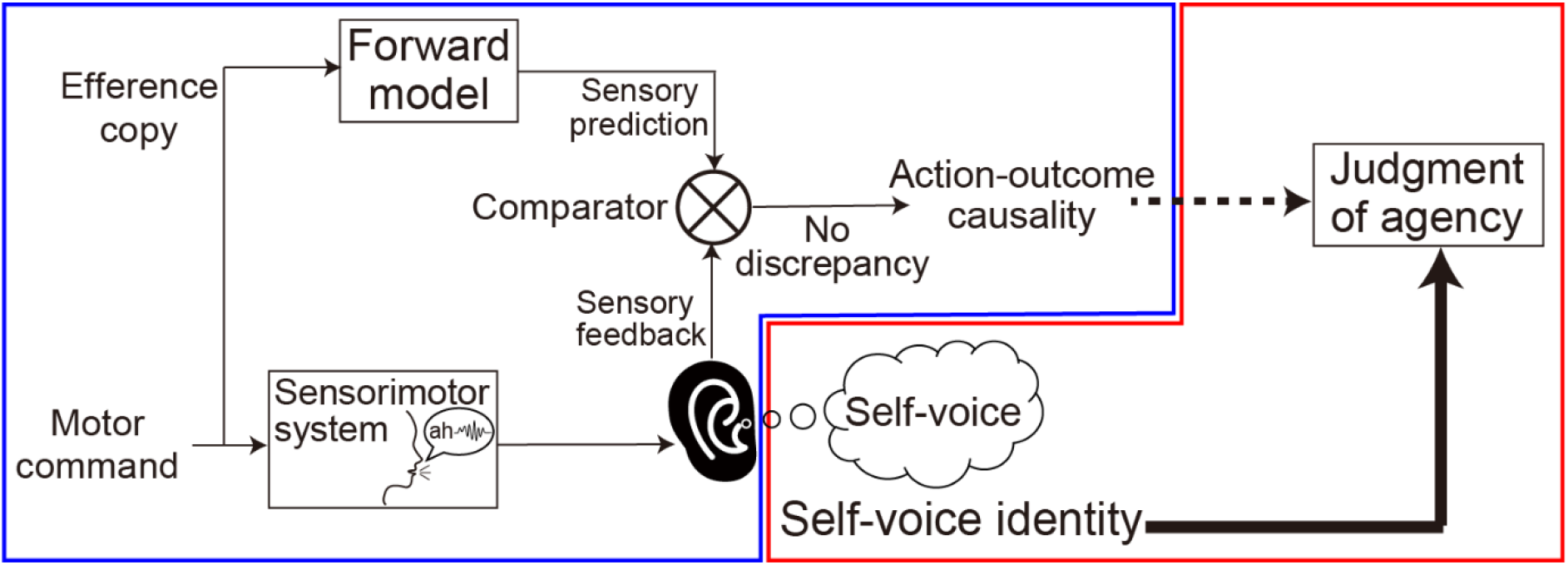
Hypothetical processes behind the sense of agency over speech. First, a sensory outcome of speech is predicted by the internal forward model. The sensory prediction is compared with actual sensory feedback. The discrepancy between them determines the estimation of causality between action and outcome (i.e., between speech and feedback voice). Sensorimotor processes are highlighted in the blue frame. By contrast, the self-voice identity (whether the feedback voice is one’s own or not) affects the judgment of agency over speech. The cognitive process shown in the red frame also constitutes the sense of agency.

In addition to action-outcome causality, another component that may affect the sense of agency over speech is self-voice identity. A sensory outcome of speech contains a sign of the self, an element of the physical self (Gillihan & Farah, 2005), in its acoustic quality. Such self-sign enables people to quickly and precisely identify whether a voice is their own or that of another. A theoretical framework suggests that the sense of agency is not simply a direct reflection of a low-level sensorimotor process of comparing an action with its outcome. Context cues, background beliefs, and post-hoc inferences (i.e., rationalizations) also affect agency judgment at the cognitive level (Synofzik et al., 2008; Synofzik et al., 2013). It is therefore plausible that the process to identify self-voice based on self-sign embedded in acoustic quality affects the judgment of agency over speech (red frame in Figure 1). Previous studies have investigated voice attribution (whether a heard voice is attributed to oneself or another) by distorting voice feedback, and they found dysfunction in patients with AHs (Goldberg et al., 1997; Johns et al., 2006; Johns & McGuire, 1999; Johns et al., 2001). Such voice attribution indeed reflects the effect of self-voice identity as well as that of action-outcome causality; however, these two effects have not been properly distinguished. Consequently, relative contribution of each effect (or that of an interaction between them) to the judgment of agency over speech remains poorly understood.

The current study investigated in detail the effects of the action-outcome causality and self-voice identity on the sense of agency over speech using both an implicit measure (without a direct self-report on the agency by participants) and an explicit measure (with a direct self-report). We first examined an action-outcome causality perceived during speech by measuring the temporal compression of a perceived interval between speech and voice feedback. This compression, termed the intentional binding effect, is often used as an implicit measure of the sense of agency (Caspar et al., 2016; Caspar et al., 2020; Haggard, 2017; Haggard et al., 2002; Moore & Obhi, 2012; Yoshie & Haggard, 2013; Yoshie & Haggard, 2017). The sensory processing for detecting action (i.e., speech) and outcome (i.e., feedback voice) timings is considered independently of the judgment of whether the feedback voice is one’s own (i.e., self-voice identity judgment). Thus, the implicit measure can access a sensorimotor process at a lower level than the cognitive process of self-voice identity. We distorted the pitch of a feedback voice and examined whether the perceived interval would be more compressed in the pitch-undistorted than in the pitch-distorted condition. Our second experiment examined the effect of self-voice identity on the judgment of agency. We required participants to explicitly report on their agency (how much they felt they had caused the voice to be heard) when we manipulated the self-voice identity by giving a pitch-distorted or undistorted self-voice following their speech as voice feedback. A time interval was inserted between their speech and the voice feedback. According to previous studies on hand/limb movement, an action-outcome temporal mismatch is critical to the loss of causality between them, which eventually causes loss of the sense of agency (Asai & Tanno, 2007; Farrer et al., 2008; Franck et al., 2001; Imaizumi & Asai, 2017; Imaizumi & Tanno, 2019; Maeda et al., 2012; Maeda et al., 2013). Thus, we manipulated the interval length to examine how self-voice identity interacts with causality in the judgment of agency. Finally, we examined an association of proneness to AH with implicit and explicit measures of agency.

## Results

### Experiment 1: intentional binding during speech

In the first experiment, we examined the sense of agency over speech in the intentional binding paradigm (Figure 2A; for details, see Methods: Experiment 1). Twenty-nine participants joined the experiment. They vocalized a vowel sound from the Japanese syllabary (“ah,” “i,” “u,” “e,” or “o”) into a microphone. Following a 200-ms, 400-ms, or 600-ms interval, they heard their voice through a headphone. The feedback voice was pitch-shifted upward or downward by seven semitones or presented without any distortion (high-/low-pitch and neutral conditions, respectively; see Supplementary Movie for a voice sample under each condition). Participants then reported an estimated interval between their speech and the voice feedback using a numeric keypad. We randomly presented the 45 conditions (three levels of speech-feedback interval × three levels of voice distortion × five levels of syllable sound).

**Figure 2.**
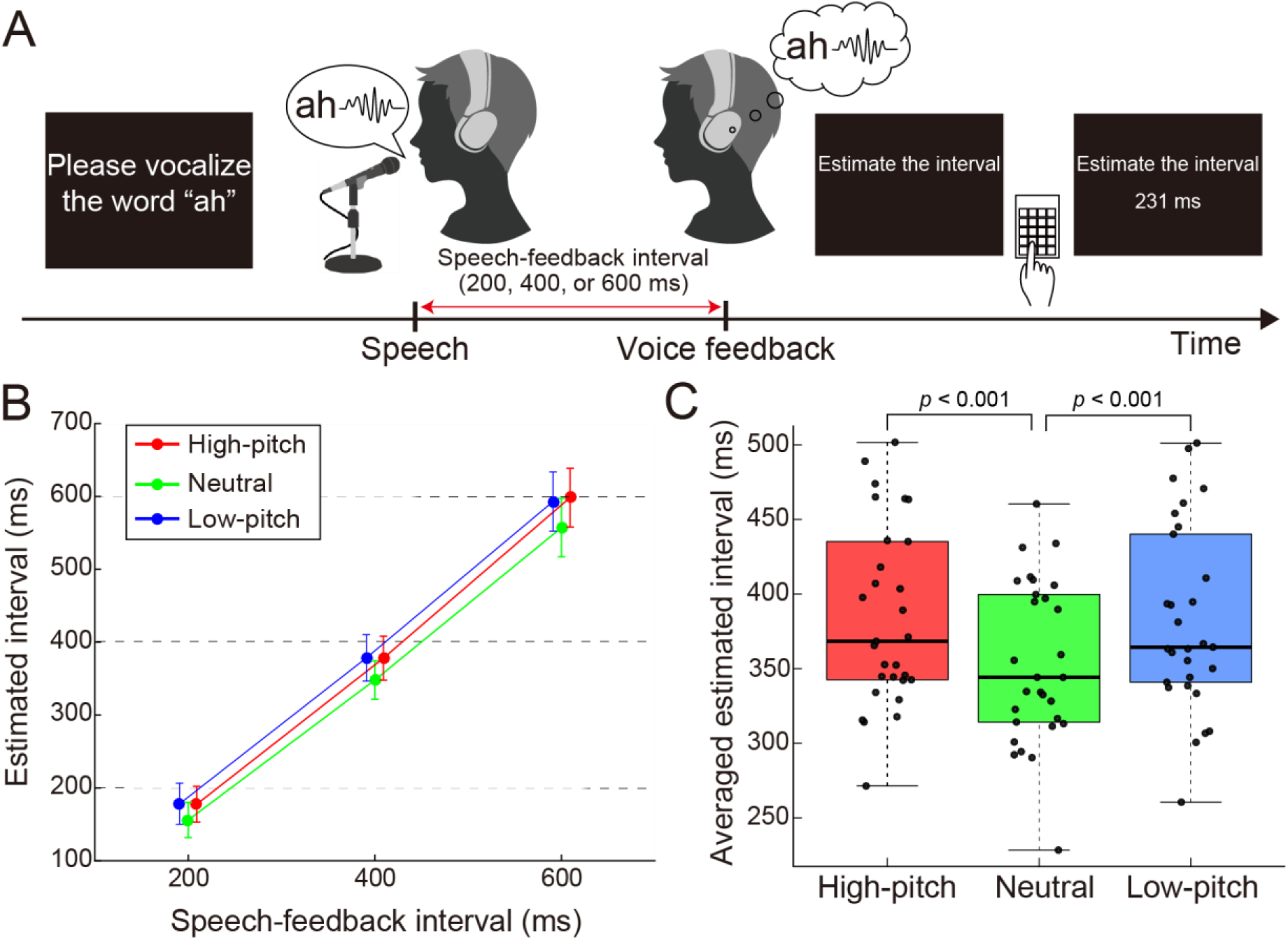
Experiment 1. (**A**) Intentional binding paradigm during speech. Participants vocalized a sound and heard their voice through a headphone following a short interval (200, 400, or 600 ms). They reported a perceived interval between their speech and voice feedback. (**B**) Mean of the estimated intervals in each pitch and speech-feedback interval. Red, green, and blue circles and lines correspond to the high-pitch, neutral, and low-pitch conditions, respectively. Error bars indicate 95% confidence intervals (CIs). Horizontal dotted lines denote the actual speech-feedback interval. Note that the circles of the high- and low-pitch conditions are shifted slightly rightward and leftward, respectively, for display purpose. (**C**) Means of the estimated intervals across the three speech-feedback intervals. In each box plot, the central horizontal line indicates the median, and the bottom and top edges correspond to the 25th and 75th percentiles. Each dot corresponds to one participant.

We analyzed estimated intervals using a 3 × 3 repeated-measures analysis of variance (ANOVA) with within-subject factors of pitch and speech-feedback interval (Figure 2B and “Data Summary” in Supplementary Data). We found a significant main effect of pitch (*F*(2, 56) = 18. 3, *p* < 0.001, *η_p_^2^* = 0.39) and a significant main effect of speech-feedback interval (*F*(1.22, 34.3) = 245.9, *p* < 0.001, *η_p_^2^* = 0.90, Greenhouse-Geisser corrected) but no significant interaction (*F*(2.74, 76.68) = 0.76, *p* = 0.51, *η_p_^2^* = 0.026, Greenhouse-Geisser corrected). Note that the effect size of the pitch (*η_p_^2^* = 0.39) is comparable to or larger than the effect sizes in the previous studies which examined participants’ estimated action-outcome intervals (Barlas et al., 2017; Caspar et al., 2015; Caspar et al., 2016; Caspar et al., 2020). Following the previous study on intentional binding (Suzuki et al., 2019), we calculated the mean of the estimated intervals across the three speech-feedback intervals to examine a simple effect of pitch (Figure 2C; see also Supplementary Figure 1 for each speech-feedback interval). We compared the means in the neutral condition with those in the high- and low-pitch conditions. The comparisons revealed that the estimated intervals in the neutral condition were shorter than those in the high-pitch (*t*(28) = 5.54,*p* < 0.001, Cohen’s *d* = 0.53, two-tailed paired *t*-test with Bonferroni correction) and low-pitch conditions (*t*(28) = 4.90, *p* < 0.001, Cohen’s *d* = 0.49, two-tailed paired *t*-test with Bonferroni correction). Collectively, the distorted voice feedback weakens the compression of the perceived interval between speech and voice feedback.

### Control experiment for Experiment 1

We were concerned that the participants would feel difficulty in detecting the onset of the distorted voice, which might cause longer estimated intervals in the high- and low-pitch conditions than those in the neutral condition. To examine this possibility, we conducted a control experiment in which the same participants in Experiment 1 reported estimated intervals between the perceived time of a beep and a prerecorded voice without online vocalization (Figure 3A). We analyzed the estimated intervals using a 3 × 3 repeated-measures ANOVA with within-subject factors of pitch and beep-voice interval (Figure 3B and “Data Summary” in Supplementary Data). As a result, we found a significant main effect of beep-voice interval (*F*(1.28, 35.97) = 434.2, *p* < 0.001, *η_p_^2^* = 0.94, Greenhouse-Geisser corrected), but no main effect of pitch was significant (*F*(2, 56) = 1.04, *p* = 0.36, *η_p_^2^* = 0.036). An interaction of these two factors was significant (*F*(4, 112) = 3.13, *p* = 0.018, *η_p_^2^* = 0.10). We found that the estimated interval at the 200-ms beep-voice interval in the high-pitch condition was significantly shorter than that in the neutral condition by a post hoc paired *t*-test with Bonferroni correction (*t*(28) = 3.23, *p* = 0.0096). However, this effect was not associated with the main result in Experiment 1 (i.e., the shorter estimated interval in the neutral condition; Figure 2B). As with the analysis of Experiment 1, we compared the mean of the estimated intervals across the three beep-voice intervals in the neutral condition with that in the high-pitch and low-pitch conditions (Figure 3C; see also Supplementary Figures. S2A, S2B, and S2C for each beep-voice interval). There was no significant difference in the mean between the high-pitch and neutral conditions (*t*(28) = 0.72, *p* = 0.48, Cohen’s *d* = 0.073, twotailed paired *t*-test) or between the low-pitch and neutral conditions (*t*(28) = 0.62, *p* = 0.54, Cohen’s *d* = 0.068, two-tailed paired *t*-test).

**Figure 3.**
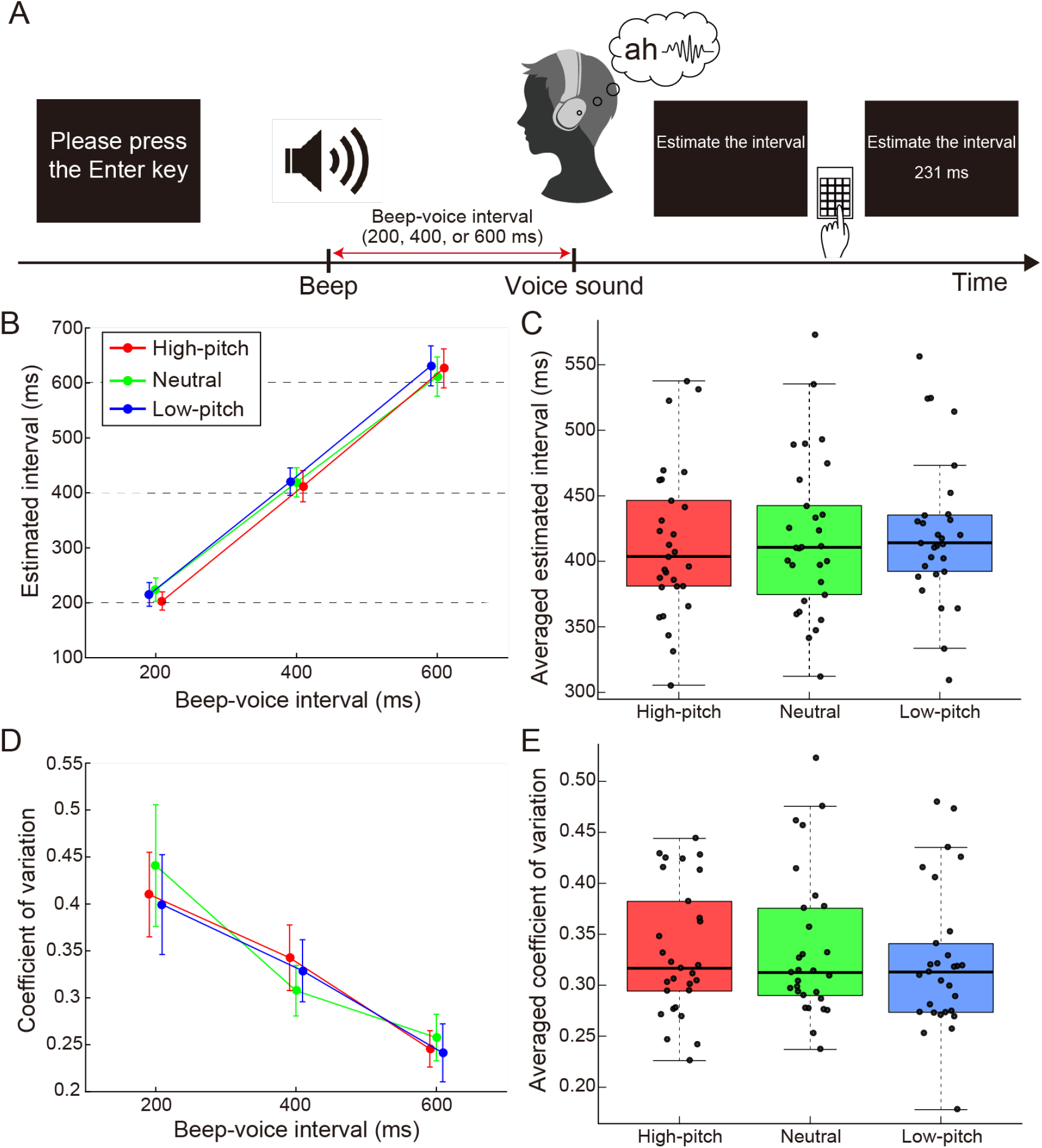
Control experiment for Experiment 1. (**A**) Experimental task. Participants heard their prerecorded voice 200, 400, or 600 ms after a beep. They reported a perceived interval between the beep and voice sound. (**B**-**E**) The mean (**B**) and coefficient of variation (**D**) of estimated intervals are shown for each pitch and beep-voice interval condition. Red, green, and blue circles and lines correspond to the high-pitch, neutral, and low-pitch conditions, respectively. Error bars indicate 95% CIs. Horizontal dotted lines in (**B**) denote the actual beep-voice interval. Note that the circles of high- and low-pitch conditions are shifted slightly rightward and leftward, respectively, for display purpose. The estimated intervals (**C**) and coefficient of variation (**E**) are averaged across the three beep-voice intervals. In each box plot, the central horizontal line indicates the median, and the bottom and top edges correspond to the 25th and 75th percentiles. Each dot corresponds to one participant.

In addition, previous experimental and computational model studies have shown that the reliability of an outcome sensory signal affects the intentional binding effect (Legaspi & Toyoizumi, 2019; Wolpe et al., 2013). To assess the reliability of auditory perception in the different pitch conditions, we calculated the coefficient of variation (CV; ratio of standard deviation to the mean) of the estimated intervals (Figure 3D and “Data Summary” in Supplementary Data). A 3 × 3 repeated measures ANOVA revealed a main effect of interval was significant (*F*(1.49, 41.8) = 52.5, *p* < 0.001, *η_p_^2^* = 0.65, Greenhouse-Geisser corrected). However, neither a main effect of pitch (*F*(1.5, 42.12) = 0.50, *p* = 0.56, *η_p_^2^* = 0.017, Greenhouse-Geisser corrected) nor an interaction (*F*(1.85, 51.94) = 1.34, *p* = 0.27, *η_p_^2^* = 0.046, Greenhouse-Geisser corrected) was significant. Moreover, we found no effect of the pitch by averaging the CVs across the three beep-voice intervals (Figure 3E; see also Supplementary Figures. S2D, S2E, and S2F for each beep-voice interval). Taken together, neither difficulty in detecting onset nor the reliability of auditory perception influenced the results in Experiment 1. Hence, the more compression found in the neutral condition than in the pitch-distorted conditions likely reflected the difference in the sense of agency. Also, although the pitch distortion might compress or expand the length of voice sound, this control experiment indicates that the changes in the length, if any, unlikely affected the difference in estimated intervals among the conditions.

### Experiment 2: subjective ratings on agency and self-voice identity

In the second experiment, we required participants to give explicit reports regarding agency over speech and self-voice evaluation (Figure 4A; for details, see Methods: Experiment 2). Twenty-eight participants who participated in Experiment 1 joined this second experiment. As with the procedure in Experiment 1, they vocalized a vowel sound in the Japanese syllabary and heard their voice through a headphone following a short interval (50, 200, 350, 500, or 650 ms). They then reported a subjective rating regarding agency (agency rating: “how much did you feel you caused the voice to be heard?”) or self-voice evaluation (self-voice rating: “how much did you feel the voice heard was your own?”) on a 9-point Likert scale from 1 (not at all) to 9 (definitely). The feedback-voice was pitch-shifted upward or downward by seven semitones or presented without distortion (high-/low-pitch and neutral conditions, respectively).

**Figure 4.**
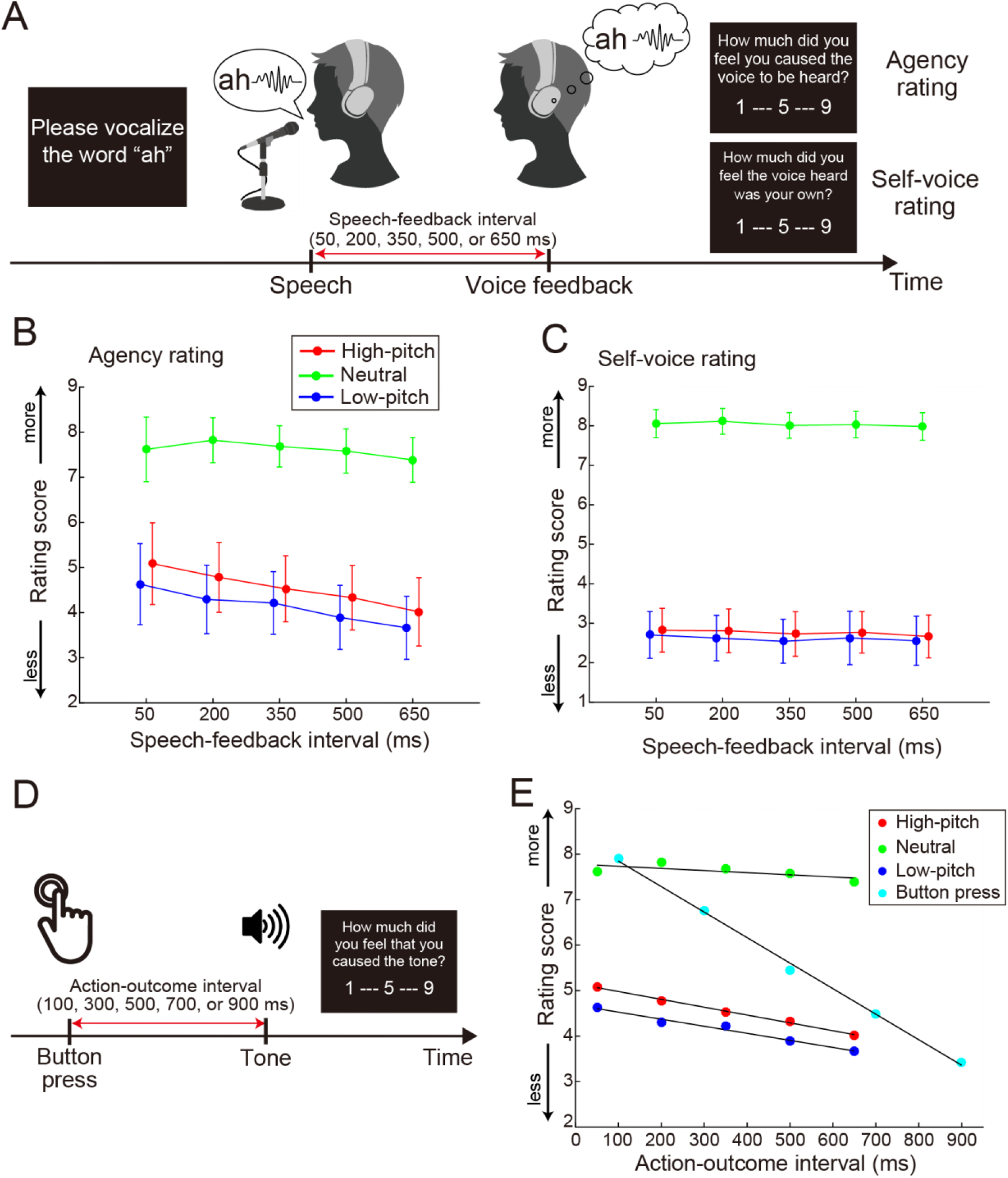
Experiment 2. (**A**) Experimental task for explicit reporting on agency and self-voice evaluation. Participants vocalized a sound and heard a feedback voice after a short interval (50, 200, 350, 500, or 650 ms). They reported how much they felt that they had caused the heard voice (agency rating) or how much they felt the heard voice was their own (self-voice rating) on a 9-point Likert scale. (**B** and **C**) Means of agency rating (**B**) and self-voice rating (**C**) in each pitch and speech-feedback interval. Red, green, and blue circles and lines correspond to high-pitch, neutral, and low-pitch conditions, respectively. Error bars indicate 95% CIs. Note that the circles of high- and low-pitch conditions are shifted slightly rightward and leftward, respectively, for display purpose. (**D**) Button-press task in Imaizumi & Tanno (2019). Participants pressed a button and heard a tone sound after a short interval (100, 300, 500, 700, or 900 ms). They reported how much they felt that they had caused the tone on a 9-point Likert scale. (**E**) Comparison of agency ratings in the speech task of the current study with those in the button-press task. The means of agency ratings are plotted as a function of action-outcome interval. Red, green, and blue circles correspond to high-pitch, neutral, and low-pitch conditions, respectively. Cyan circle denotes the mean of agency ratings in the button-press task. Black lines indicate regression lines, which were fitted to the means of ratings in the respective conditions.

Figure 4B shows the mean score of agency rating in each pitch and speech-feedback interval (see also “Data Summary” in Supplementary Data). We conducted a 3 × 5 repeated-measures ANOVA with within-subject factors of pitch and speech-feedback interval. As a result, a main effect of pitch was significant (*F*(1.39, 37.49) = 79.3, *p* < 0.001, *η_p_^2^* = 0.75, Greenhouse-Geisser corrected), but no main effect of speech-feedback interval was significant (*F*(1.15, 30.96) = 2.67, *p* = 0.11, *η_p_^2^* = 0.090, Greenhouse-Geisser corrected). In addition, an interaction between these two factors was significant (*F*(4.85, 130.98) = 5.10,*p* < 0.001, *η_p_^2^* = 0.16, Greenhouse-Geisser corrected). Post-hoc tests revealed a significant simple main effect of speech-feedback interval in the high-pitch condition (*F*(1.32, 35.6) = 4.29, *p* = 0.036, *η_p_^2^* = 0.14, Greenhouse-Geisser corrected) and a marginal effect in the low-pitch condition (*F*(1.29, 34.95) = 3.27, *p* = 0.069, *η_p_^2^* = 0.11, Greenhouse-Geisser corrected), but no significant effect was found in the neutral condition (*F*(1.28, 34.52) = 0.80, *p* = 0.40, *η_p_^2^* = 0.029, Greenhouse-Geisser corrected).

To our surprise, we did not find a significant effect of the speech-feedback interval, at least for the neutral condition, despite the fact that a temporal mismatch between an action and its outcome is critical for a decrease in the sense of agency in hand/limb movement tasks (Asai & Tanno, 2007; Farrer et al., 2008; Franck et al., 2001; Imaizumi & Asai, 2017; Imaizumi & Tanno, 2019; Maeda et al., 2012; Maeda et al., 2013). Next, we compared the results using our speech paradigm with those using the button-press paradigm reported by Imaizumi and Tanno (2019). In this previous study (Figure 4D), participants pressed a button followed by a short interval (100, 300, 500, 700, or 900 ms) and heard a tone via a headphone. The participants then rated the perceived sense of agency over the tone by answering the question: “How much did you feel that you had caused the tone?” We fitted a linear regression model to each of 34 participants’ rating score as a function of action-outcome interval (ms). We also fitted the model to the agency ratings in Experiment 2. As a result, the mean of the slopes was −5.6 (SD: 2.1) in the button-press task, whereas the means were −0.47 (3.6), −1.7 (3.9), and −1.6 (4.1) in the neutral, high-pitch, and low-pitch conditions of the speech task, respectively (Figure 4E). The slope in the button-press task was significantly and negatively steeper than the values under any condition of the speech task (high-pitch condition: *t*(60) = 4.99,*p* < 0.01, Cohen’s *d* = 1.27; neutral condition: *t*(60) = 7.00,*p* < 0.01, Cohen’s *d* = 1.79; and low-pitch condition: *t*(60) = 5.00,*p* < 0.01, Cohen’s *d* = 1.28, two-tailed *t*-test). Furthermore, we compared the results using our speech paradigm with those from the two different tasks using hand movement (Asai & Tanno, 2007; Imaizumi & Asai, 2017). Note that participants in these studies reported agency judgment differently from those in Imaizumi and Tanno (2019) (i.e., 9-point Likert scale versus two-alternative forced-choice). Thus, the procedure in our study was more similar to that in Imaizumi and Tanno (2019) than those in the two hand-movement tasks. We found the slopes in the hand-movement tasks were significantly and negatively steeper than the values in at least the neutral condition of the speech task (for details, see Supplementary Texts: 1. Comparison of agency judgment between hand/limb movement and speech tasks and Supplementary Figure 3). These comparisons clarified that the judgment of agency over speech was more immune to action-outcome mismatch than that of agency over hand/limb movement.

Next, regarding self-voice rating (Figure 4C and “Data Summary” in Supplementary Data), we found a significant main effect of pitch (*F*(1.56, 42.16) = 274.9, *p* < 0.001, *η_p_^2^* = 0.91) but neither a significant main effect of speech-feedback interval (*F*(1.5, 40.45) = 0.94, *p* = 0.37, *η_p_^2^* = 0.034) nor a significant interaction (*F*(4.73, 127.58) = 0.19, *p* = 0.96, *η_p_^2^* = 0.0070). Thus, both agency rating and self-voice rating were sensitive to the pitch distortion of the feedback voice, but the self-voice rating was not significantly affected by the speech-feedback interval.

### Association with proneness to having auditory hallucinations

Finally, we examined whether the indices measured in Experiments 1 and 2 (estimated intervals, agency ratings, and self-voice ratings in the three pitch-conditions) could explain the individual difference in proneness to having AHs, as quantified by a questionnaire (Auditory Hallucination Experience Scale 17: AHES-17). First, we compared the AHES-17 scores collected in the present study *(N* = 28) with a score distribution of a large sample *(N* = 827) in the previous study (Asai et al., 2011b). The scores in the present study ranged from 27 to 65 (mean: 43.5, SD: 11.2). This score range fell within the mean ± 2SD of the large-sample distribution (mean: 48.2, SD: 11.7; Figure 5A). Thus, the participants in our study did not show extreme proneness to AHs.

**Figure 5.**
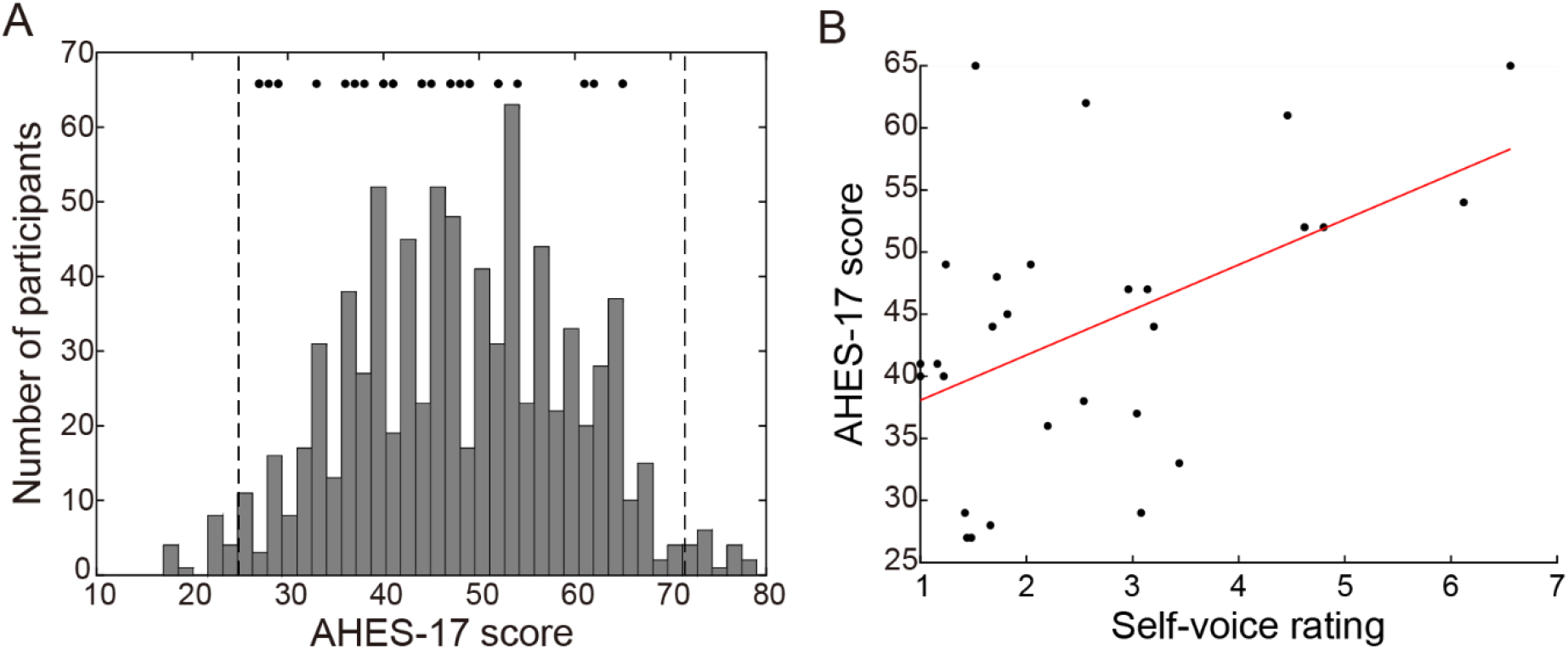
Association with proneness to auditory hallucinations. (**A**) Distribution of AHES-17 scores. The histogram shows the distribution of 827 participants’ scores collected in Asai et al. (2011b). Dots above the histogram indicate 28 participants’ data in the current study. Broken vertical lines indicate the mean ± 2SD of the distribution of the 827 samples. (B) Correlation between self-voice rating in low-pitch condition and AHES-17 scores. Correlation coefficient was 0.49 (*t*(26) = 2.87, *p* = 0.0080). Each dot corresponds to one participant. The red line denotes the regression line.

Next, we conducted a stepwise multiple regression analysis, using MATLAB function *stepwisefit*, to add or remove indices according to how their inclusion affects the model. The indices that significantly improved the model (*F* statistic, *p* < 0.05) were added to the model as a predictor. As a result, the self-voice ratings in the low-pitch condition alone could significantly explain the AHES-17 scores (*R*^2^ = 0.24, *F*(1,26) = 8.24, *p* = 0.0080). Figure 5B shows the correlation between the self-voice ratings in the low-pitch condition and AHES-17 scores. The correlation coefficient was significantly larger than zero according to a permutation test *(r* = 0.49, *t*(26) = 2.87, *p* = 0.0080, 95% confidence interval (CI) = [0.14, 0.73]). Four participants scored over 60, relatively higher among the 28 participants in the current study. We also checked the correlation excluding the four participants from the data, but the correlation was still significantly larger than zero (*r* = 0.46, *t*(22) = 2.43, *p* = 0.024, 95% CI = [0.070, 0.73]). Thus, the above results indicate that individual differences in proneness to AH were associated with the process of evaluating self-identity in hearing a voice.

## Discussion

The current study examined the two possible components that affect the sense of agency over speech: action-outcome causality and self-voice identity. In the first experiment, we found more compression of the estimated intervals between speech and voice feedback when the participants heard an undistorted self-voice than a distorted one (Figures 2B and 2C). This compression, corresponding to an intentional binding effect, indicates that stronger action-outcome causality can be perceived in hearing self-voice following one’s speech. In the second experiment, we investigated the effect of self-voice identity, manipulated by pitch distortion, on the judgment (rating) of agency. We first found that self-voice identity strongly affects the agency rating (Figure 4B) and the self-voice rating (Figure 4C). We next found that the agency rating over speech was immune to a temporal mismatch between action and its outcome (i.e., between speech and feedback voice), especially in the case where self-voice identity was preserved (i.e., when undistorted self-voice was fed back) (Figure 4E). This property was not confirmed in the judgment of agency over hand/limb movement in previous studies. Finally, we found an association of proneness to AH with the self-voice rating but not with either an implicit or explicit measure of agency (Figure 5B).

In Experiment 1, we demonstrated, for the first time to our knowledge, the intentional binding effect in a naturalistic speech paradigm (but see Limerick et al. (2015), which used a speech interface controlled by voice command). What caused the lesser compression in the pitch-distorted conditions than in the neutral condition? A recent Bayesian model of the sense of agency rationally explains the intentional binding effect. It posits three requirements for confidence in causal estimation, which is how the model theorizes the sense of agency (Legaspi & Toyoizumi, 2019): (1) temporal consistency in the action-outcome interval, (2) prior belief that the action caused the outcome, and (3) reliability of sensorimotor signals informing action and outcome timings. In the current experiment, we randomly presented three speech-feedback intervals and three pitch-distorted conditions on a trial-by-trial basis. Therefore, regarding the first two of the above requirements, temporal consistency in the action-outcome effect and prior causal belief are comparable in all pitch-distorted conditions. Thus, referring to the computational model, a plausible cause of the lesser compression in the pitch-distorted conditions is having unreliable sensorimotor signals of action and outcome timings. As suggested by the comparator model (Blakemore, et al., 2000b; Frith et al., 2000a), the sensorimotor system predicts a sensory outcome of speech (i.e., feedback voice) and then compares the predicted and actual outcomes (Ford, 2016; Jones & Fernyhough, 2007; Seal et al., 2004). People already establish an internal forward model of speech in the course of daily activities. As the established internal model predicts non-distorted voice as an outcome of speech, the pitch distortion produces a discrepancy in acoustic quality between the predicted and actual outcomes. The neural activity involved in the sensorimotor processing may be perturbed by this discrepancy, thus reducing the sensory reliability of action and outcome timings and estimating weakened causality between speech and voice feedback. This explanation is consistent with the perturbed neural activity in hearing distorted voice feedback found in previous neurophysiological studies (Ford, 2016; Heinks-Maldonado et al., 2005; Heinks-Maldonado et al., 2007). Accordingly, such a low-level sensorimotor process plausibly lessened the compression in the pitch-distorted conditions.

Experiment 2 investigated the effect of self-voice identity on the sense of agency over speech at the cognitive judgment level. The first finding in this experiment was the large effect of self-voice identity on agency over speech. The pitch distortion remarkably decreased the agency rating scores (Figure 4B) as well as the self-voice rating (Figure 4C), and these decreases were almost comparable. This effect appeared in the vertical discrepancies between the rating scores in the neutral condition and those in the two distorted conditions in Figures. 4B and 4C. These results suggest that the sign of the self in the acoustic quality of feedback voice strongly affects the judgment of agency over speech.

The second finding in Experiment 2 was a small effect of speech-feedback interval on the agency rating. A temporal mismatch between an action and its outcome is generally a critical factor in reducing the sense of agency (Asai & Tanno, 2007; Farrer et al., 2008; Franck et al., 2001; Imaizumi & Asai, 2017; Imaizumi & Tanno, 2019; Maeda et al., 2012). However, compared with the button-press paradigm (Imaizumi & Tanno, 2019), we found that the agency ratings less steeply decreased as a function of action-outcome interval in all pitch conditions of this speech paradigm (Figure 4E). The critical difference between the button-press paradigm and our speech paradigm is whether the outcome of an action (i.e., tone or voice sound, respectively) contains a sign of the self. In our additional experiment (see Supplementary Texts: 2. Self-voice identification of prerecorded voice as a function of pitch distortion), we confirmed that 1) the participants recognized the undistorted voice as their own and 2) the level of pitch distortion in our experiment (i.e., ±7 semitones) was adequate for them to recognize the distorted voice as a non-self-voice. These results suggest that participants linked the feedback voice with their speech once they recognized the voice as their own, whereas this link was disconnected once they recognized the voice as someone else’s. This top-down cognitive effect of self-voice identity was so dominant that the agency judgment became immune to action-outcome temporal relation that could contribute to the agency judgment as a bottom-up process. Thus, our findings highlighted the dominant top-down effect of self-voice identity on the judgment of agency over speech (red frame in Figure 1). In the future experiment, one’s face can be used as another type of the self-sign that causes a top-down effect on the agency over speech. That is, the judgment of agency over speech may also be immune to action-outcome temporal relation if people recognize a self-sign in a visualized face during speech. Such an experiment needs a feedback voice consistent with one’s own or someone else’s face movement presented on a monitor. Further studies are needed to examine other types of a sign of the self and their effects on the sense of agency over speech by incorporating “enfacement illusions” (Sforza et al., 2010; Tsakiris, 2008) or virtual reality techniques (Banakou & Slater, 2014) into the experimental paradigm in the current study.

The third finding in Experiment 2 was the significant interaction between pitch and action-outcome interval shown by the ANOVA of the agency rating scores. The post-hoc tests for individual pitch-conditions revealed that the simple main effect of the interval was significant only in the pitch-shifted conditions (red and blue lines in Figure 4B), but not in the neutral condition (green line in Figure 4B). We can explain the reasons for this interaction in terms of the dominance of self-voice identity as follows. On the one hand, a non-distorted self-voice was heard following a speech in the neutral condition. The participants might perceive this phenomenon as reasonable based on their daily experience. Thus, self-voice identity dominantly affected the agency rating in this condition, resulting in the rating being immune to action-outcome temporal mismatch. On the other hand, it is unreasonable to hear a distorted self-voice following a speech presented in the pitch-distortion conditions. Thus, the dominance of the self-voice identity was relatively weaker in those conditions than in the neutral condition. Therefore, a mixed effect of self-voice identity and action-outcome temporal mismatch determined the agency ratings.

In addition to the above findings, we found that neither implicit nor explicit measures of the sense of agency but instead self-voice rating in the low-pitch condition could explain the individual differences in proneness to AHs (Figure 5B). In addition to the low-pitch condition, we found a moderate correlation between the self-voice ratings in the high-pitch condition and AHES-17 scores *(r* = 0.32, *t*(26) = 1.85, *p* = 0.094, 95% CI = [-0.058, 0.62]). By contrast, the correlations with the self-voice ratings in the neutral condition and with the other indices (i.e., estimated intervals and agency ratings) scored at most *r* = 0.10. Note that we give a detailed discussion of the positive correlation between the self-voice ratings and AHES-17 scores in our study (Figure 5B), which seems to be the opposite tendency of previous studies, in Supplementary Texts (3. Relationship between self-voice identity and proneness to auditory hallucination). These results suggest that the cognitive process of evaluating self-voice identity in a feedback voice is critical to explaining the AH-like experiences in (non-psychotic) participants. One possible reason why only the self-voice ratings were associated with proneness to AHs is because the scale we evaluated in this study more strongly reflect proneness to dissociative than that to schizophrenic AHs. Although AHs are indeed a representative symptom of schizophrenia, clinical observations have shown that patients with dissociative identity disorder also exhibit first-rank symptoms of schizophrenia, including AHs (Kluft, 1987; Ross et al., 1990). Unlike pathological dissociation, even some healthy individuals can experience mild dissociation, such as talking with imaginary friends inside the head. Such mild dissociation might moderately disrupt self-voice identity. In addition, correlations with two factors of AHES-17 scores support our above interpretation. The factor analysis in the previous study revealed two factors that influence AHES-17 scores (Asai et al., 2011a). The first factor includes AH-like experience caused by internal speech/thinking and related to music or others’ voices, which is common with dissociative and schizophrenic AHs. By contrast, the second factor is associated with delusions or thought insertion, which is the representative positive symptoms of schizophrenia. Asai et al. (2011a) classified questionnaire items into two groups according to relative loading of the two factors. We calculated correlation coefficients between the self-voice ratings in the low-pitch condition and the total score within the first group (internal speech/thought) or the second group (delusions or thought insertion) while removing variance explained by the other factor from the two variables (i.e., partial correlation). As a result, the correlation coefficient with the scores of the first group (*r* = 0.31) was larger than that with the scores of the second group (*r* = 0.11). Thus, we can assume that the AHES-17 scores in this study strongly reflected dissociative AH-like experiences such as internal speech/thought, music, and other’s voice. Taken together, the significant correlation between the scores and self-voice ratings might be reasonable if this assumption is valid. To validate this assumption, we need further studies that compare self-voice ratings of participants showing schizotypal traits with those showing dissociative traits.

In the present study, there is a limitation on bone conduction’s influence upon the sense of agency over speech. Voices heard through our ears are not the only sensory feedback in speech. Bone conduction is also one of the primary sensory inputs active while we are speaking. We cannot deny bone conduction’s effect on the sense of agency measured in the present task, although this effect was reduced by mixing white noise into the feedback voice (see Methods: Apparatus). We have not examined how much participants could hear their voice directly (i.e., not through a headphone), either. However, the bone conduction’s effect should not be different across all the pitch conditions. Therefore, bone conduction unlikely affected our main conclusion that distorted self-voice feedback attenuated the sense of agency over speech.

In sum, this study has characterized the mechanism underlying the sense of agency over speech. The first experiment indicated that the predictive sensorimotor system enabled the brain to estimate causality between an action and its outcome during speech (blue frame in Figure 1). This low-level bottom-up process has also been characterized as the primary component of the sense of agency over hand/limb movement. The second experiment, by contrast, highlighted the uniqueness of the sense of agency over speech; that is, the sign of the self in the acoustic quality of voice (i.e., self-voice identity) constitutes a crucial component of the sense of agency at the cognitive judgment level (red frame in Figure 1). This top-down effect was so dominant that the judgment is immune to a result processed by the low-level sensorimotor system. These findings shed light on the nature of the subjective experience of self-agency during speech. They would also be informative for a deeper understanding of aberrant experience in AHs.

## Methods

### Participants

Twenty-nine healthy volunteers (15 males and 14 females) with a mean age of 21.7 (19-25 years of age) participated in Experiment 1 and the control experiment for Experiment 1. The same volunteers, except for one female (i.e., 28 volunteers), participated in Experiment 2. We determined the sample sizes based on our preliminary experiment (see Supplementary Texts: 4. Preliminary experiment on the effect of intentional binding during speech). For the sample size calculation, we performed a power analysis for repeated measures ANOVA using G*power 3.1 with power selected at 0.95, effect size (*η_p_^2^*) at 0.42, and alpha at 0.05 (Faul et al., 2007). The experimental protocol was approved by the ethics committee at the University of Tokyo. Written informed consent was obtained from all volunteers in accordance with the latest version of the Declaration of Helsinki.

### Apparatus

All experiments were conducted inside a soundproof room. Participants seated themselves in front of a microphone (SENNHEISER MD42) on a desk and wore a headphone (HyperX Cloud Revolver Pro Gaming Headset, Kingston Technology Company). The microphone was connected to an effector (BEHRINGER VIRTUALIZER Pro) via an amplifier (AT-MA2 MICROPHONE AMPLIFIER, Audio-Technica). We used the effector to change the pitch of a spoken voice and to insert a time interval between speech and voice feedback. Note that the effector’s sample rate was so high (46 kHz) that people could not detect any lag due to the experiment’s pitch distortion. White noise was mixed into the feedback voice using a sound mixer (YAMAHA MW10C) to reduce direct voice transmission (not via the effector and headphone) and bone conduction. A custom Python program running on a laptop computer placed on a desk presented the visual stimuli and collected the participants’ responses.

### Questionnaire on proneness to auditory hallucination

Before the participants performed the task of Experiment 1, they answered a questionnaire on their proneness to auditory hallucination (AH). We used the Auditory Hallucination Experience Scale 17 (AHES-17), which was translated into Japanese (Asai et al., 2011a) and had been used in previous studies (Asai et al., 2011b; Asai & Tanno, 2013; Sugimori et al., 2011). The AHES-17 is a short version of the AHES and includes 17 self-reporting questions, each of which participants answer on a 5-point Likert scale. The range of possible scores is from 17 to 85.

### Experiment 1: Intentional binding during speech

#### Experimental task and procedure

Participants were required to report the perceived interval between their speech and voice feedback (Figure 2A). At the beginning of each trial, a message was presented on the screen to instruct them to utter one of five vowel sounds in the Japanese syllabary (“ah,” “i,” “u,” “e,” or “o”). Participants pressed the enter key when they were ready, and the message on the screen disappeared. Then, they vocalized the instructed sound into the microphone. Following a short interval (200, 400, or 600 ms), participants heard the spoken sound through a headphone. The feedback voice was pitch-shifted upward or downward by seven semitones or presented without any distortion (high- or low-pitch or neutral condition, respectively). After the message “Estimate the interval” was presented on the screen, they reported the estimated interval between their speech and voice feedback using a numeric keypad. They were instructed to vocalize a sound as briefly and clearly as possible, and they were required to speak closely to the microphone. Participants reported the estimated interval in three digits, ranging from 100 to 999 ms. They performed four trials for each of the 45 conditions (three levels of speech-feedback interval × three levels of voice distortion × five levels of sound) in random order (i.e., 180 trials in total).

Before the main task session, participants performed a practice session to get accustomed to the estimation of an interval between speech and voice feedback. They vocalized the sound instructed on the screen and heard their spoken voice, as in the main task. Then, they reported the estimated interval between their speech and voice feedback. They were informed of the actual interval as a feedback on their estimation accuracy. In the practice session, we set a speech-feedback interval at 11 different levels from 0 to 1000 ms with a 100-ms interval. The feedback voice was not distorted (i.e., the neutral condition only). Participants conducted 22 trials in total (2 trials × 11 conditions in random order).

#### Data analysis

We averaged estimated intervals across the five sounds in each pitch and speech-feedback interval condition for each participant. Then, we performed a 3 × 3 repeated-measures ANOVA on the estimated intervals with within-subject factors of pitch (high, neutral, and low) and speech-feedback interval (200, 400, and 600 ms). If the sphericity assumption for ANOVA was violated, we applied Greenhouse-Geisser correction.

### Control experiment for Experiment 1

#### Experimental task and procedure

Participants were required to report the interval between the perceived time of a beep and a prerecorded voice sound (Figure 3A). At the beginning of a trial, a message was presented on the screen prompting participants to press the enter key. A beep sound was presented after a random interval (2,000 to 5,000 ms) from the keypress. Following a short interval (200, 400, or 600 ms), a recorded voice was replayed through a headphone. Then, the participants reported the estimated interval between the time of a beep and voice sound using a numeric keypad. They were instructed to report the estimated interval in three digits, ranging from 100 to 999 ms. Before the experimental task, we recorded each participant’s voice as they vocalized five vowel sounds in the Japanese syllabary (“ah,” “i,” “u,” “e,” or “o”). We extracted vocalization sections from the recorded auditory file. We distorted the pitch by shifting it upward and downward by seven semitones (high- and low-pitch condition, respectively) using audio software (Audacity, https://www.audacityteam.org/). Participants conducted four trials for each of the 45 conditions (three levels of beep-voice interval × three levels of voice distortion × five levels of sound) in random order (i.e., 180 trials in total). They performed a practice session before the main task session. In the practice session, the beep-voice interval was set from 0 to 1000 ms with 100-ms intervals. As the practice session in Experiment 1, they received the actual interval as a feedback after reporting their estimated interval. The recorded voice was replayed without pitch distortion. Participants performed two trials for each of the 11 conditions in random order (i.e., 22 trials in total).

#### Data analysis

We averaged estimated intervals across the five sounds in each pitch and beep-voice interval condition for each participant. We performed a 3 × 3 repeated-measures ANOVA on estimated intervals with within-subject factors of pitch (high, neutral, and low) and beep-voice interval (200, 400, and 600 ms). If the sphericity assumption for ANOVA was violated, we applied Greenhouse-Geisser correction. In the post-hoc test for the ANOVA, Bonferroni correction was used for multiple comparisons.

### Experiment 2: Subjective report on agency over speech and self-voice evaluation

#### Experimental task and procedure

Participants were required to report subjective ratings regarding agency over speech and self-voice identification (Figure 4A). At the beginning of each trial, a message was presented on the screen to instruct them to vocalize one out of five sounds in the Japanese syllabary (“ah,” “i,” “u,” “e,” or “o”). The message disappeared immediately after the keypress, and participants spoke the instructed sound into the microphone. Following a short interval (50, 200, 350, 500, or 650 ms), the spoken sound was heard through the headphone. The feedback-voice was pitch-shifted upward or downward by seven semitones or presented without distortion (high- or low-pitch or neutral condition, respectively). After a message was presented on the screen, they reported a subjective rating regarding agency over speech (agency rating) or regarding self-voice evaluation (self-voice rating). The message was “how much did you feel you had caused the voice to be heard?” for the agency rating or “how much did you feel the voice heard was your own?” for the self-voice rating. The rating was scored on a 9- point Likert scale from 1 (not at all) to 9 (definitely). As in Experiment 1, we instructed participants to vocalize the sound as briefly and clearly as possible and to speak closely to the microphone.

In Experiment 2, we did not require participants to perform any practice session. We were concerned about the possibility that they might confuse the agency rating with the self-voice rating if they reported both in a single trial or either in random order. Therefore, we required them to report agency ratings in the first half block and then self-voice ratings in the last half block. In each block, they conducted two trials for each of 75 conditions (five levels of speech-feedback interval × three levels of voice distortion × five levels of sound) in random order. Together with the two blocks, they conducted 300 trials in total.

#### Data analysis

We averaged rating scores across the five sounds in each pitch and speech-feedback interval condition for each participant. We performed a 3 × 5 repeated-measures ANOVA on rating scores with within-subject factors of pitch (high, neutral, and low) and speech-feedback interval (50, 200, 350, 500, and 650 ms). If the sphericity assumption for ANOVA was violated, we applied Greenhouse-Geisser correction.

## Data availability

Data for each participant from Experiment 1, Control experiment for Experiment 1, and Experiment 2 are available in a spreadsheet file of Supplementary Data. For details, see README sheet of the file.

## Acknowledgements

H.I. was supported by JSPS KAKENHI Grant Numbers 18H01098, 19H05725, and 19H01777. T.A. was supported by JSPS KAKENHI Grant Number 17K13971. H.I. and T.A. were also supported by “Research and development of technology for enhancing functional recovery of elderly and disabled people based on noninvasive brain imaging and robotic assistive devices,” Commissioned Research of National Institute of Information and Communications Technology (NICT) and Japan Agency for Medical Research and Development (AMED) (grant JP18dm0307008). The authors are grateful to Dr Takaki Maeda (Keio University School of Medicine) for his insightful comments on this manuscript from the viewpoint of a psychiatrist. We would like to thank Hiroki Tarumi for his assistance with data collection and Ayuko Misu and Marina Sano for their helpful discussions on the experimental design.

## Author contribution statement

R.O., T.A., and H.I. designed the study; R.O. collected the data in all of the experiments; T.A. and S.I. summarized the data of the previous studies, which were compared with the data in the current study. R.O. analyzed the data; R.O. and H.I. wrote the manuscript; T.A. and S.I. reviewed and approved the final version of the manuscript.

## Supplementary Information

**Supplementary Texts**

1. Comparison of agency judgment between hand/limb movement and speech tasks
2. Self-voice identification of prerecorded voice as a function of pitch distortion
3. Relationship between self-voice identity and proneness to auditory hallucination
4. Preliminary experiment on the effect of intentional binding during speech

**Supplementary Figures**

**Supplementary Figure 1** Estimated intervals for speech-feedback intervals in Experiment 1.

**Supplementary Figure 2** Estimated interval and coefficient of variation in control experiment.

**Supplementary Figure 3** Comparison with results of hand/limb movement tasks.

**Supplementary Figure 4** Results of self-voice identification task.

**Supplementary Figure 5** Difference in voice-distortion level between the previous and present studies and its effect on the self-voice rating.

**Supplementary Figure 6** Interval estimations in preliminary experiment.

**Supplementary Text**

### 1. Comparison of agency judgment between hand/limb movement and speech tasks

We investigated whether the explicit judgment of agency in our speech paradigm was more immune to action-outcome mismatch than that in hand/limb movement paradigms. We compared the results in the two other studies in addition to those in Imaizumi and Tanno (2019), which were compared with our results in the main text. The experimental paradigm we first investigated was the mouse-control task conducted by Asai and Tanno (2007). Eleven participants were required to control a mouse device to reach one of the four corners of the screen from the starting point (Supplementary Figure 3A). The cursor on the screen moved correspondingly to the mouse movement with a time delay (0, 250, 500, 750, or 1000 ms). They then judged whether they felt that they had moved the cursor on their own (i.e., two-alternative forced choice). We fitted a linear regression model to the “yes” response ratio of each of 11 participants as a function of time delay (ms). We also fitted the model to the agency ratings, reported on a 9-point Likert scale, in Experiment 2. Note that the rating scores were divided by 9 to compare the regression slopes fitted to the Likert-scaled rating in our study with the response ratio of the agency judgment in the previous study. As a result, the mean of the slopes was −0.75 (SD: 0.39) in the mouse-control task, whereas the means were −0.052 (0.40), −0.19 (0.44), and −0.17 (0.46) in the neutral, high-pitch, and low-pitch conditions in the speech task, respectively (Supplementary Figure 3B). The slope in the mouse-control task was significantly and negatively steeper than those in every condition of the speech task (high-pitch condition: *t*(37) = 3.68,*p* < 0.01, Cohen’s *d* = 1.31; neutral condition: *t*(37) = 4.90,*p* < 0.01, Cohen’s *d* = 1.74; and low-pitch condition: *t*(37) = 3.64,*p* < 0.01, Cohen’s *d* = 1.30, two-tailed *t*-test).

The second experimental paradigm we investigated was the hand-movement task conducted by Imaizumi and Asai (2017). In the study (Supplementary Figure 3C), 16 participants placed their left-hand with palm down and moved the hand to form the target hand position instructed prior to the movement. The video feedback was presented on the screen with a time delay (50, 250, 500, 1000, or 1500 ms). Participants then judged whether they felt as if they were controlling the movement of the hand on the screen on their own (i.e., two-alternative forced choice). As with the above comparison, we fitted a linear regression model to the “yes” response ratio of each participant as a function of time delay (ms). As a result, the mean of the slopes was −0.33 (SD: 0.29) in the hand-movement task (Supplementary Figure 3D). We found a marginal difference between the slope in the hand-movement task and that in the neutral condition of our speech task (*t*(42) = 2.44, *p* = 0.056, Cohen’s *d* = 0.77, two-tailed *t*-test with Bonferroni correction), while the slopes in the high- and low-pitch conditions were not significantly different (high-pitch condition: *t*(42) = 1.15, *p* = 0.26, Cohen’s *d* = 0.36; and low-pitch condition: *t*(42) = 1.25, *p* = 0.22, Cohen’s *d* = 0.39, two-tailed *t*-test).

### 2. Self-voice identification of prerecorded voice as a function of pitch distortion

#### Methods

##### Participants

Twenty-six healthy volunteers with a mean age of 21.7 years (19-25 years old) participated in the experiment (14 males and 12 females). They also participated in Experiments 1 and 2, except for one male and two females. Note that we excluded two participants’ data from the analysis because the estimated point of subjective equalities (PSEs) showed an inappropriate value (lower than zero, see below).

##### Experimental task and procedure

Participants started a trial by pressing the enter key. A prerecorded voice sound was presented after a random interval (2,000 to 5,000 ms) from the keypress. They judged whether the sound was self-voice or non-self-voice by pressing a button on a numeric keypad. Before the experimental task, we recorded each participant’s voice while they were vocalizing five vowel sounds of the Japanese syllabary (“ah,” “i,” “u,” “e,” or “o”). We extracted vocalization sections from the recorded file. Pitch was distorted into 15 levels from −7 to +7 semitones with a one-semitone interval using audio software (Audacity). Note that positive and negative values denote upward and downward pitch-shift, respectively. Participants conducted two trials for each of the 75 conditions (15 levels of voice alteration × five levels of sound) in random order (i.e., 150 trials in total).

##### Data analysis

We calculated the ratio of “self-voice” response by dividing the number of trials in which participants judged “self-voice” by the total number of trials in each pitch condition. We divided all of the data into two sub-datasets: upward-shift (from 0 to +7 semitone) and downward-shift (from −7 to 0 semitone). A sigmoid function was fitted to each dataset using a Matlab (MathWorks, Natick, MA, USA) *glmfit* function. We estimated two PSEs for each participant and tested the difference between them using a two-tailed paired *t*-test.

#### Results

Supplementary Figure 4 shows the ratio of the “self-voice” response at each pitch. The ratio declined as the amount of pitch-shift increased and reached approximately zero at the ±7-semitone shift. Note that the voices were distorted by the ±7-semitone shift in Experiments 1 and 2. We also found that the PSE of the upward-shift data was significantly smaller than that of the downward-shift data (*t*(23) = 4.34, *p* < 0.001, Cohen’s *d* = 0.91, two-tailed paired *t*-test).

### 3. Relationship between self-voice identity and proneness to auditory hallucination

We found that those who showed high AH proneness were more likely to identify the pitch-distorted voice as their own than those who showed low AH proneness (i.e., positive correlation between self-voice ratings and AHES-17 scores in Figure 5B). However, earlier studies on verbal self-monitoring in AH patients showed the opposite tendency: AH patients were more likely to judge distorted self-voice as non-self-voice than healthy controls(Goldberg et al., 1997; Johns et al., 2006; Johns & McGuire, 1999; Johns et al., 2001). Clues to understanding this seemingly inconsistency are the difference in distortion level between the previous and current studies and participants’ discriminability in self-other attribution of sensory signals as follows (Supplementary Figure 5). Asai (2016) found a low discriminability of self-other attribution in schizotypal participants. In his experiment, participants were asked participants to move a cursor on a screen using a pen-tablet device while he parametrically included a control signal from a pen of another participant into the cursor movement. He measured individual differences in the discriminability of self-other attribution of the cursor movement (i.e., how sensitively participants discriminate the observed movement originated from the self or other). As a result, higher schizotypal participants showed low discriminability of self-other attribution. Although the current task is different from that in Asai (2016), we assume that AH patients or people with high AH proneness have poorer discriminability of self-other voice than healthy controls or people with low AH proneness. In the present study, the pitch of feedback-voice was much distorted (by seven semitones). Thus, the participants with low AH proneness, who had high discriminability, were more likely to report distorted voice as non-self-one (blue solid circle in the yellow shaded area of Supplementary Figure 5). By contrast, the participants with high AH proneness had relatively low discriminability and showed the tendency to recognize the distorted voice as their own (red solid circle in the yellow shaded area of Supplementary Figure 5). In the previous studies(Goldberg et al., 1997; Johns et al., 2006; Johns & McGuire, 1999; Johns et al., 2001), the pitch was slightly distorted. Therefore, due to low discriminability, patients with AHs were more likely to recognize the distorted voice as other’s (red dotted circle in the green shaded area of Supplementary Figure 5) in the condition where healthy controls recognized feedback-voice as their voice (blue dotted circle in the green shaded area of Supplementary Figure 5). Thus, our results are, from the perspective of the discriminability of self-other voice, consistent with the previous findings.

### 4. Preliminary experiment on the effect of intentional binding during speech

Before the main experiment, we conducted a preliminary experiment to assess the effect of intentional binding during speech. Twenty-four participants, who were different from the participants of the main experiments, joined the preliminary experiment. The experimental task was the same as that in Experiment 1, but feedback voice was either distorted by shifting the pitch upward by seven semitones or not distorted (high-pitch and neutral conditions, respectively). We averaged the estimated intervals between speech and voice feedback across participants (Supplementary Figure. 6). Note that the estimated intervals were averaged across the five-vowel sounds. We performed a 2 × 3 repeated-measures ANOVA on estimated intervals with within-subject factors of pitch (high-pitch and neutral) and speech-feedback interval (200, 400, and 600 ms). We found a significant main effect of pitch *(F(1*, 23) = 16.68,*p* < 0.001, *η_p_^2^* = 0.42) and a significant main effect of speech-feedback interval (*F*(1.36, 31.17) = 373.31, *p* < 0.001, *η_p_^2^* = 0.94, Greenhouse-Geisser corrected) but no significant interaction (*F*(1.94, 44.59) = 2.14, *p* = 0.13, *η_p_^2^* = 0.085, Greenhouse-Geisser corrected).

## Supplementary Figures

**Supplementary Figure 1.**
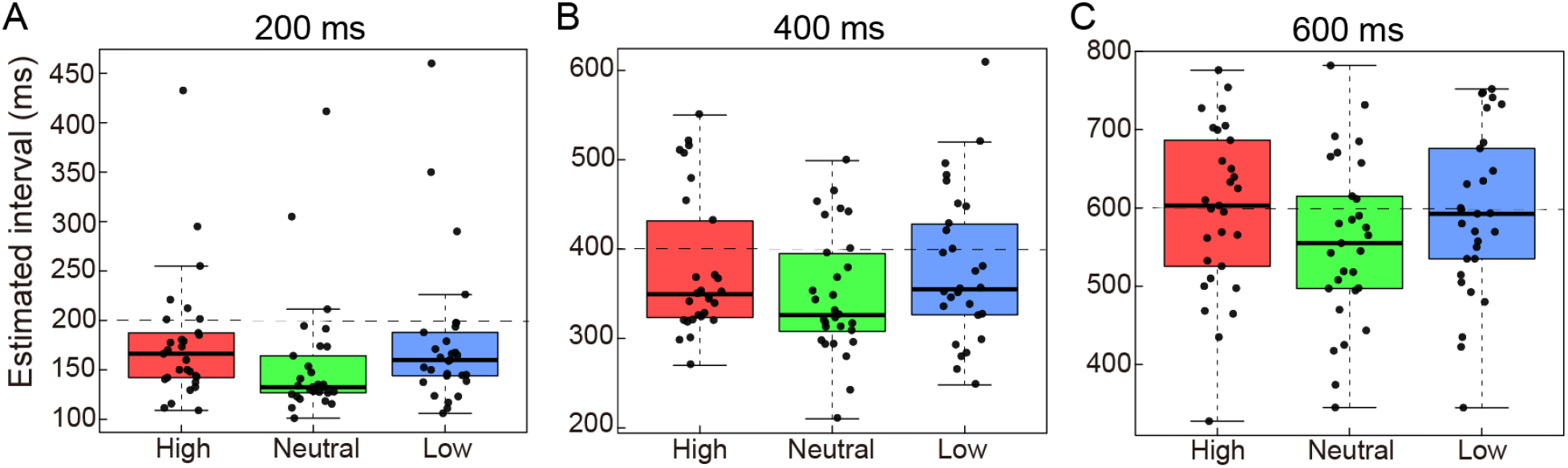
Estimated speech-feedback intervals in Experiment 1. In the 200-ms interval condition (**A**), the estimated intervals in the neutral condition were shorter than those in the high-pitch (*t*(28) = 4.38, *p* < 0.001, Cohen’s *d* = 0.34, two-tailed paired *t*-test with Bonferroni correction) and low-pitch conditions (*t*(28) = 5.13,*p* < 0.001, Cohen’s *d* = 0.29). In the 400-ms interval condition (**B**), the estimated intervals in the neutral condition were shorter than those in the other two conditions (high-pitch condition: *t*(28) = 4.41,*p* < 0.001, Cohen’s *d* = 0.39; low-pitch condition: *t*(28) = 4.01,*p* < 0.001, Cohen’s *d* = 0.37). In the 600-ms interval condition (**C**), the estimated intervals in the neutral condition were shorter than those in in the other two conditions (high-pitch condition: *t*(28) = 4.41, *p* = 0.0048, Cohen’s *d* = 0.39; low-pitch condition: *t*(28) = 4.01, *p* = 0.024, Cohen’s *d* = 0.34). Horizontal dotted lines indicate actual speech-feedback intervals. In each box plot, the central horizontal line indicates the median, and the bottom and top edges correspond to the 25th and 75th percentiles. Each dot corresponds to one participant.

**Supplementary Figure 2.**
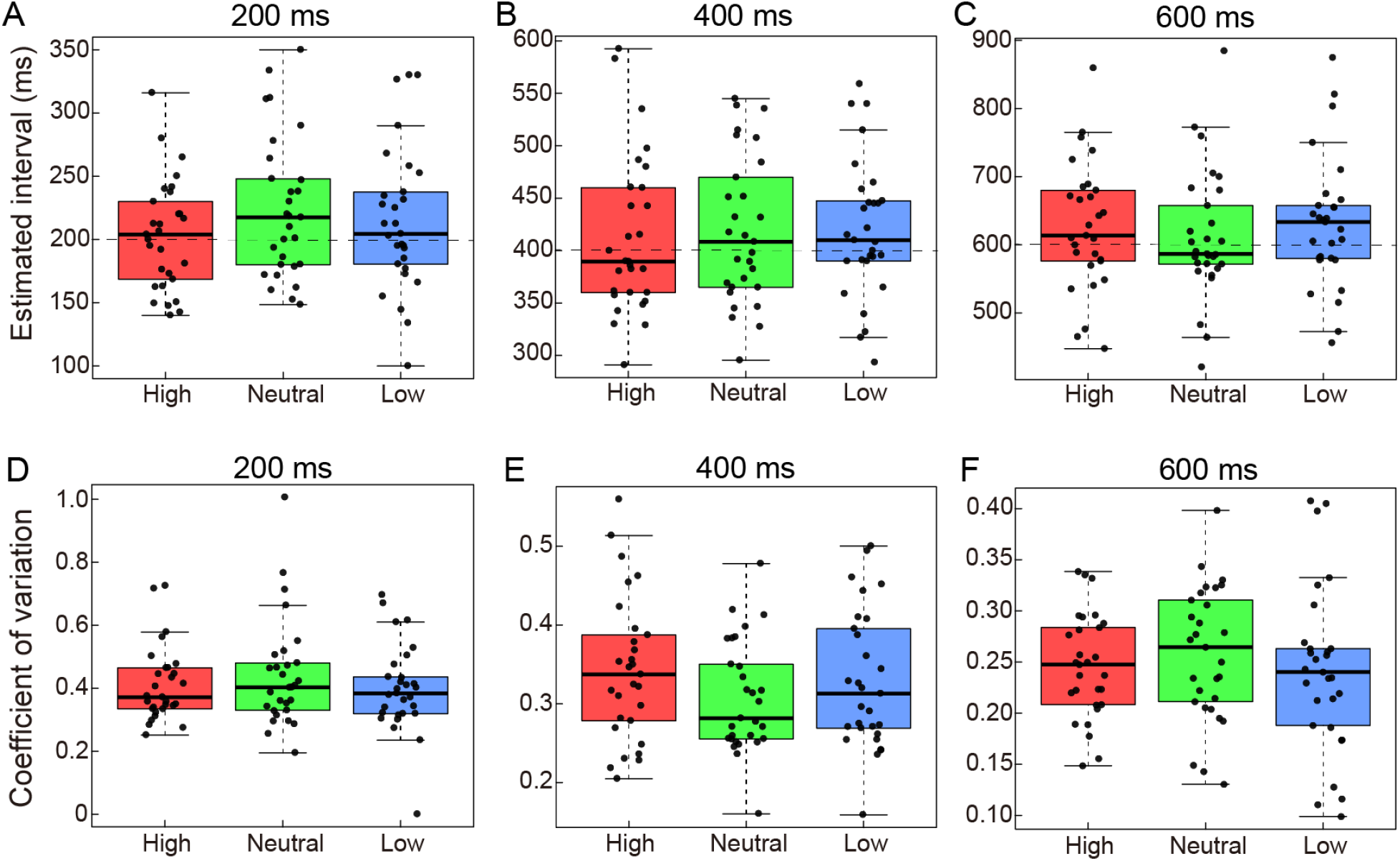
Estimated interval and coefficient of variation in Control Experiment. (**A**-**C**) Estimated intervals for 200-ms (**A**), 400-ms (**B**), and 600-ms (**C**) beep-voice intervals. Horizontal dotted lines indicate actual beep-voice intervals. (**D**-**F**) Coefficient of variations for 200-ms (**D**), 400-ms (**E**), and 600-ms (**F**) beep-voice intervals. In each box plot, the central horizontal line indicates the median, and the bottom and top edges correspond to the 25th and 75th percentiles. Each dot corresponds to one participant.

**Supplementary Figure 3.**
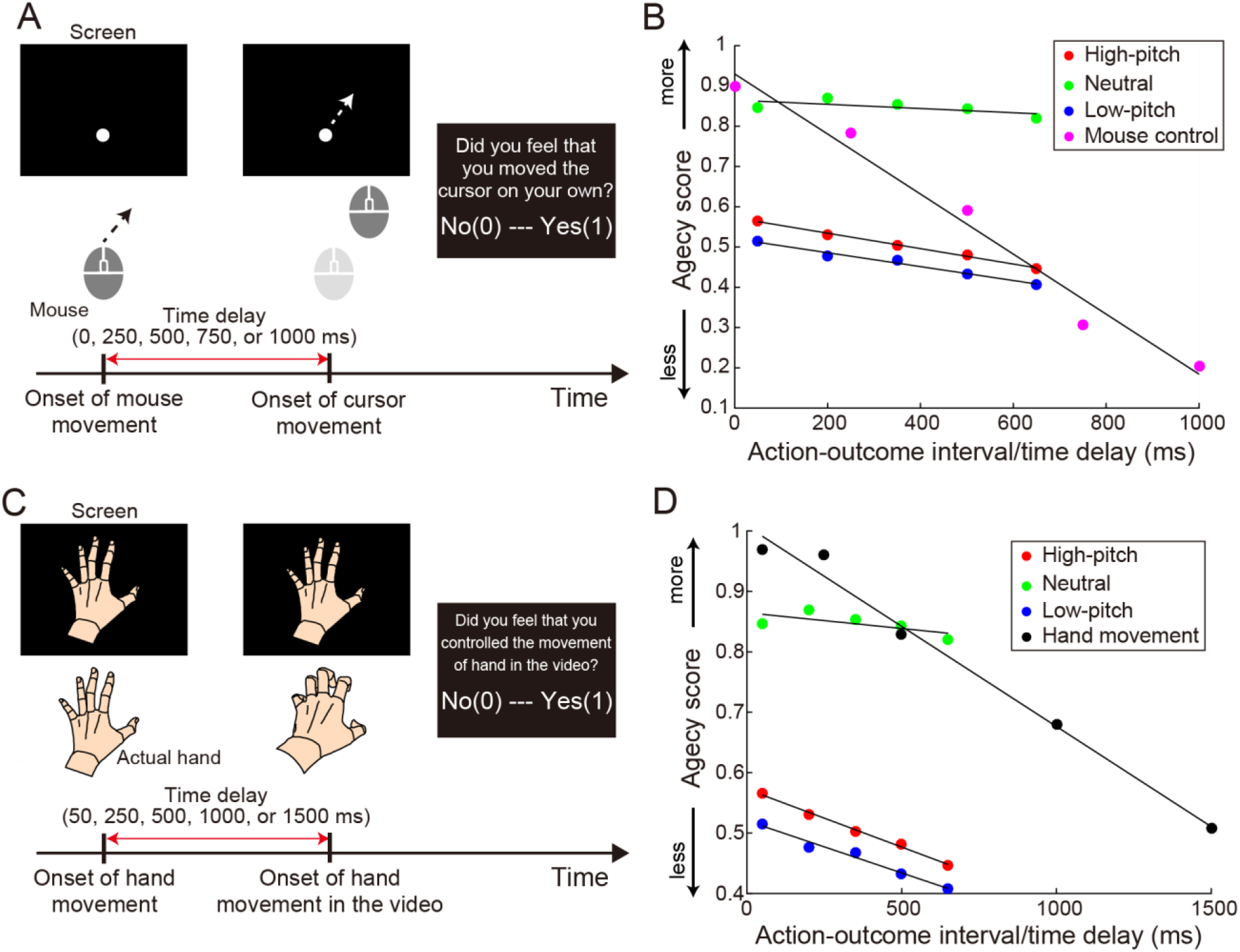
Comparison of results of hand/limb movement tasks. (**A**) Mouse-control task in Asai and Tanno (2007). Participants controlled a mouse device to move a cursor on the screen. The cursor movement was delayed relative to the mouse movement (0, 250, 500, 750, or 1000 ms). They judged whether they felt that they had moved the cursor on their own. (**B**) Comparison of agency judgment in the speech task of the current study with that in the mouse-control task. The means of agency ratings are plotted as a function of action-outcome interval while the means of “yes” response ratio are plotted as a function of time delay. Note that the agency ratings were reported on a 9-point Likert scale in the current study; consequently, they were divided by 9 to compare regression slopes fitted to the rating scores with that fitted to the response ratio in the mouse-control task. Red, green, and blue circles correspond to high-pitch, neutral, and low-pitch conditions, respectively. Magenta circles denote the means of the “yes” response ratio in the mouse-control task. Black lines indicate regression lines, which were fitted to the mean scores in the respective conditions. (**C**) Hand-movement task in Imaizumi and Asai (2017). Participants moved their hand to form an instructed posture. Video feedback on a screen was presented with a time delay (50, 250, 500, 1000, or 1500 ms). They judged whether they felt as if they were controlling the movement of the hand on the screen on their own. (**D**) Comparison of agency judgment in the speech task with that in the hand-movement task. The means of agency ratings are plotted as a function of action-outcome interval while the means of “yes” response ratio are plotted as a function of time delay. Note that the agency ratings in our study were divided by 9. Red, green, and blue circles correspond to high-pitch, neutral, and low-pitch conditions, respectively. Black circles denote the means of the “yes” response ratio in the hand-movement task. Black lines indicate regression lines, which were fitted to the mean scores in the respective conditions.

**Supplementary Figure 4.**
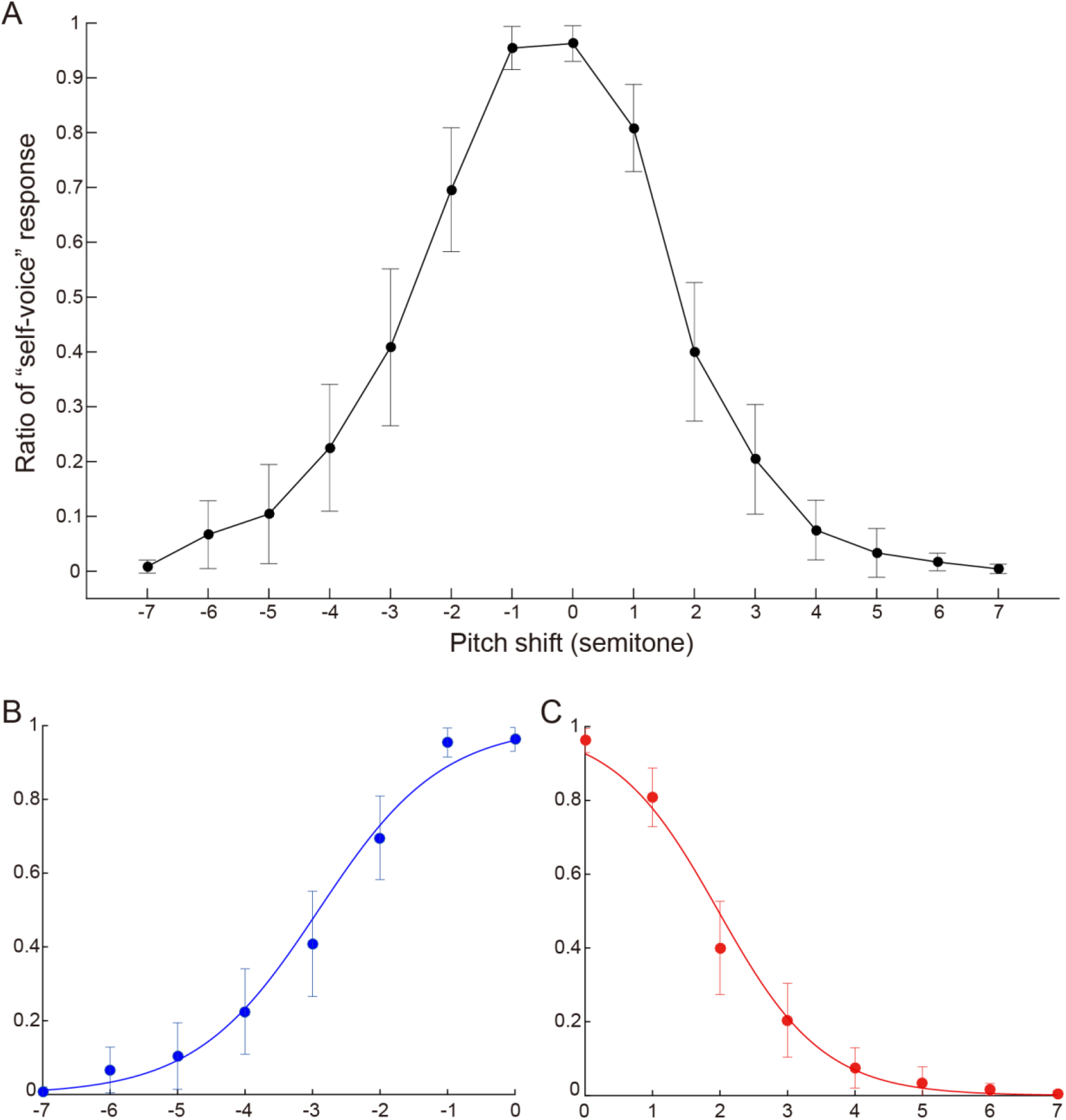
Results of self-voice identification task. (**A**) Ratio of “self” response in each pitch. A sigmoid function was fitted separately to data in the downward (**B**) and upward shift conditions (**C**). The blue and red curves denote the fitting results. Error bars indicate 95% confidence intervals (CIs).

**Supplementary Figure 5.**
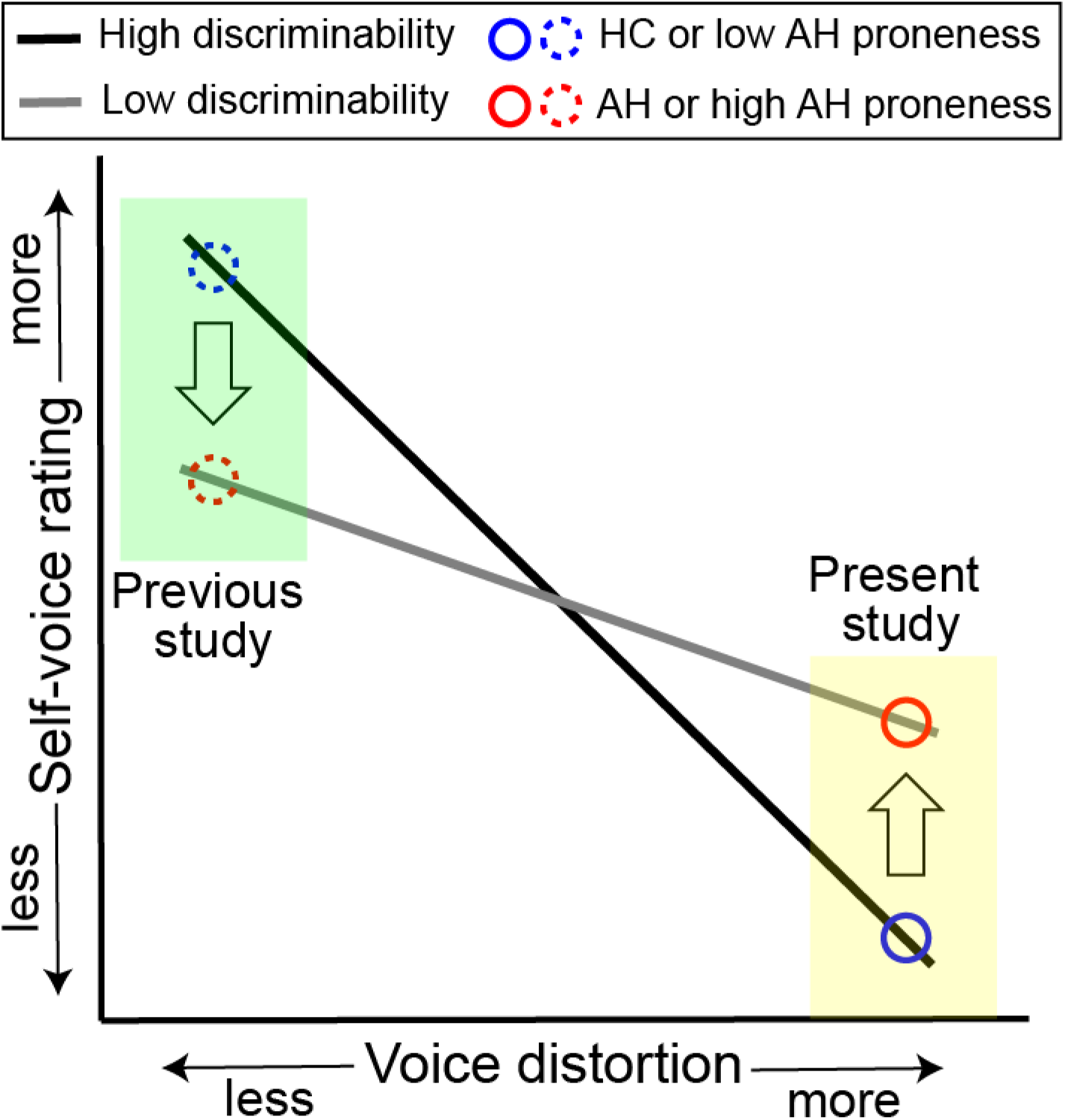
Difference in voice-distortion level between the previous and present studies and its effect on the self-voice rating. Lines indicate self-voice rating scores as a function of the level of voice-distortion. A steep black line represents a high discriminability of self-other voice, while a shallow gray line represents a low discriminability. Participants with a low discriminability (high auditory hallucination [AH] proneness in the present study) are more likely to recognize a much-distorted voice as self-voice (a red solid circle in the yellow area). However, participants with a low discriminability (patients with AH in the previous studies) are more likely to recognize a less distorted voice as non-self-voice (red dotted circle in the green area). Solid and dotted blue circles indicate people with low AH proneness in the present study and healthy controls (HC) in the previous studies, respectively.

**Supplementary Figure 6.**
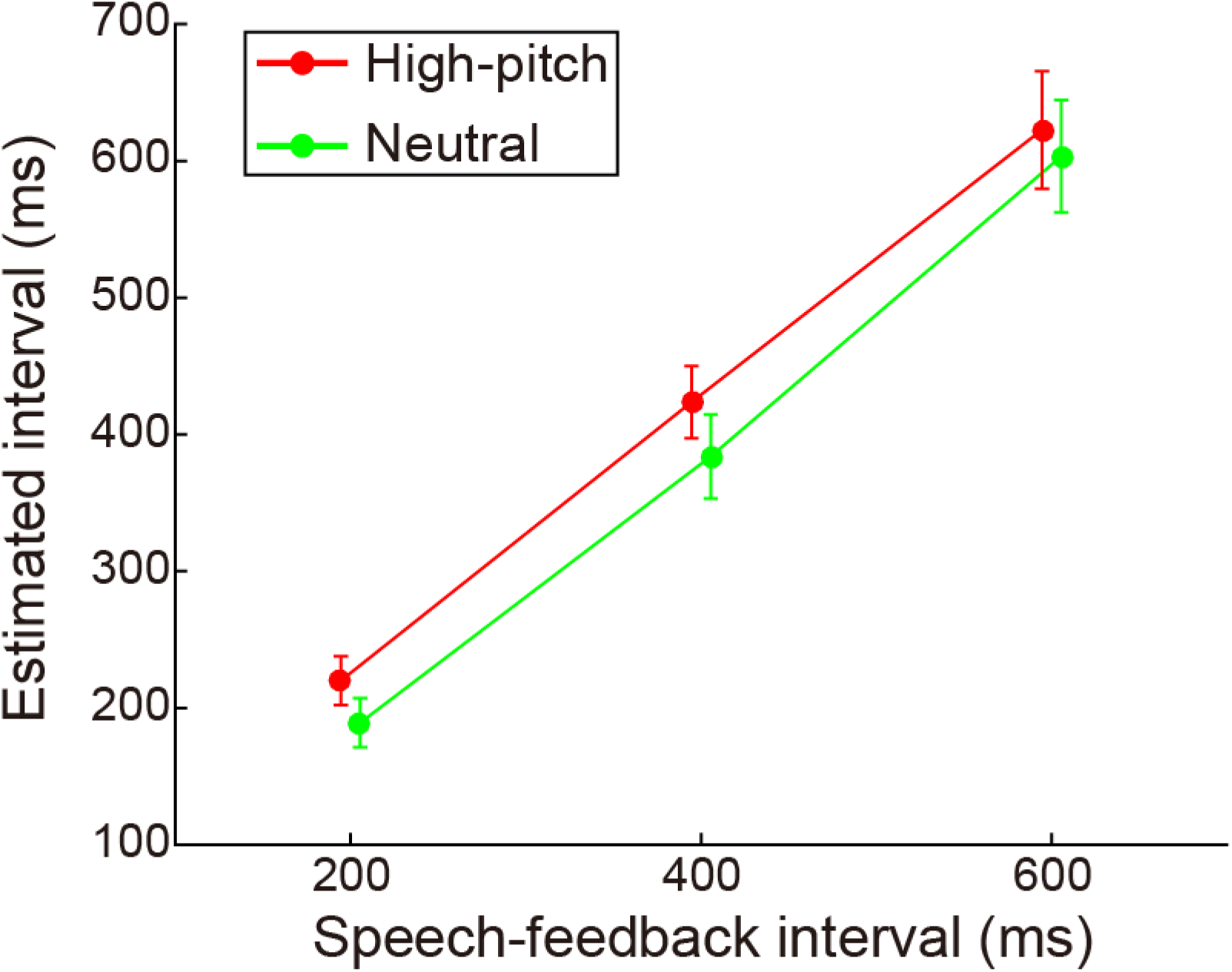
Interval estimations in the preliminary experiment. The mean of the estimated interval is plotted as a function of speech-feedback interval separately for each pitch condition. Red and green circles and lines correspond to high-pitch and neutral conditions, respectively. Error bars indicate 95% confidence intervals (CIs).

## Supplementary Movie

Examples of feedback voices in high-pitch, neutral, and low-pitch conditions. The sound is the Japanese vowel “ah.” Color panels denote sound spectrograms. The voice in the neutral condition is replayed twice in the movie. Each replay is followed by a distorted voice in either the high-pitch or low-pitch condition.

